# Classification of platelet aggregates by agonist type

**DOI:** 10.1101/805408

**Authors:** Yuqi Zhou, Atsushi Yasumoto, Cheng Lei, Chun-Jung Huang, Hirofumi Kobayashi, Yunzhao Wu, Sheng Yan, Chia-Wei Sun, Yutaka Yatomi, Keisuke Goda

## Abstract

Platelets are anucleate cells in blood whose principal function is to stop bleeding by forming aggregates for hemostatic reactions. In addition to their participation in physiological hemostasis, platelet aggregates are also involved in pathological thrombosis and play an important role in inflammation, atherosclerosis, and cancer metastasis. The aggregation of platelets is elicited by various agonists, but these platelet aggregates have long been considered indistinguishable and impossible to classify. Here we present an intelligent method for classifying them by agonist type. It is based on a convolutional neural network trained by high-throughput imaging flow cytometry of blood cells to identify and differentiate subtle yet appreciable morphological features of platelet aggregates activated by different types of agonists. The method is a powerful tool for studying the underlying mechanism of platelet aggregation and is expected to open a window on an entirely new class of clinical diagnostics, pharmacometrics, and therapeutics.

## INTRODUCTION

Platelets are non-nucleated cells in blood whose principal function is to stop bleeding by forming aggregates for hemostatic reactions (Michelson, 2012; George, 2000; Michelson, 2003; Harrison, 2005). In addition to their participation in physiological hemostasis (Michelson, 2012; George, 2000; Michelson, 2003; Harrison, 2005), platelet aggregates are also involved in pathological thrombosis (Davì and Patrono, 2007; Ruggeri, 2002). Moreover, it is known that a range of diseases or medical conditions, such as inflammation, atherosclerosis, and cancer metastasis, are closely associated with platelet aggregation (Lievens and von Hundelshausen, 2011; Engelmann and Massberg, 2013; Franco et al., 2015; Gay and Felding-Habermann, 2011). Here, the aggregation of platelets is elicited by a variety of agonists, which bind to and activate specific receptors expressed on the platelet. This leads to platelet activation and structural and functional changes of glycoprotein IIb/IIIa expressed on the platelet surface. The activated form of the glycoprotein can bind with fibrinogen, enabling platelets to interact with each other and form aggregates (Michelson, 2012; George, 2000; Michelson, 2003; Harrison, 2005; Moser et al., 2008). Despite the existence of diverse agonist types, platelet aggregates look morphologically similar and have long been thought indistinguishable since the discovery of platelet aggregates in the 19th century (Michelson, 2012; George, 2000; Michelson, 2003; Harrison, 2005). This is because morphological characteristics of platelet aggregates on a large statistical scale have been overlooked as microscopy (a high-content, but low-throughput tool) has been the only method to examine them (Finsterbusch et al., 2018; Nitta et al., 2018).

In this Report, we present an intelligent method for classifying platelet aggregates by agonist type. This is enabled by performing high-throughput imaging flow cytometry of numerous blood cells, training a convolutional neural network (CNN) with the image data, and using the CNN to identify and differentiate subtle yet appreciable morphological features of platelet aggregates activated by different types of agonists. Our finding that platelet aggregates can be classified by agonist type through their morphology is unprecedented as it has never been reported previously. The information about the driving factors behind the formation of platelet aggregates is expected to lead to a better understanding of the underlying mechanism of platelet aggregation and open a window on an entirely new class of clinical diagnostics, pharmacometrics, and therapeutics.

## RESULTS

### Development of the iPAC

Our procedure for developing an intelligent platelet aggregate classifier (iPAC) is schematically shown in Figure 1A. First, a blood sample obtained from a healthy person was separated into several different portions, into which different types of agonists were added to activate platelets while no agonist was added to the last portion for negative control (Figure S1; Transparent Methods). Here, adenosine diphosphate (ADP), collagen, thrombin receptor activator peptide-6 (TRAP-6), and U46619 were used since they are commonly used in platelet aggregation tests (Michelson, 2012; George, 2000; Michelson, 2003; Harrison, 2005). Initially, the concentrations of the agonists were carefully chosen (20 μM for ADP, 10 μg/mL for collagen, 13 μM for TRAP-6, 14 μM for U46619) to minimize variations in aggregate size between the different blood sample portions. These images were acquired through six experimental trials (Figure S2) to mitigate potential bias in the dataset that may have come from experimental variations (e.g., signal-to-noise ratio, fluctuations in optical alignment, hydrodynamic cell focusing conditions, sample preparation). Then, four different concentrations of each agonist (2, 5, 10, 20 μM for ADP, 1, 5, 10, 20 μg/mL for collagen, 1, 5, 13, 20 μM for TRAP-6, 2.8, 5.6, 14, 28 μM for U46619) were used for platelet activation to examine the potential influence of agonist concentrations on the ability to differentiate platelet aggregates by agonist type, where the concentrations were chosen by referring to the concentrations of agonists used in light transmission aggregometry and *in vitro* flow-cytometric platelet aggregation tests (Koltai et al., 2017; Granja et al., 2015). The platelet aggregates were enriched by density-gradient centrifugation to remove erythrocytes from the blood sample portions. To prevent the platelet aggregates from dissolving during imaging flow cytometry, 2% paraformaldehyde was added to the blood sample portions to fix them. In addition to this sample preparation procedure, we tested other procedures such as pipetting, vortexing, fixation, and non-fixation and identified the current procedure to be advantageous over the others in preserving the morphology of platelet aggregates (Figure S3; Transparent Methods). Second, an optofluidic time-stretch microscope (Goda et al., 2009; Jiang et al., 2017; Lei et al., 2018; Lau et al., 2016) was employed for high-throughput, blur-free, bright-field image acquisition of events (e.g., single platelets, platelet-platelet aggregates, platelet-leukocyte aggregates, single leukocytes, cell debris, remaining erythrocytes) in each sample portion (Figures S4 and S5; Transparent Methods). Third, the acquired images of the events were used to train two CNN models that classified the platelets based on their morphological features by agonist type (Figure 1B). Specifically, we first trained a CNN model with images of platelet aggregates activated by certain concentrations of agonists (12,000 images per agonist type) in order to examine their morphological changes while minimizing a potential influence of concentration-dependent factors on the morphology of the platelet aggregates. Then, we trained the other CNN model with a dataset in which the images of platelet aggregates activated by different concentrations of the agonists were equally mixed (12,000 images in total per agonist type) in order to show that different concentrations of the agonists do not perturb the CNN model’s ability to classify platelet aggregates. We employed the CNN (Krizhevsky et al., 2012) with an encoder-decoder architecture to disregard insignificant features such as background noise and keep important features in the bottleneck layer and trained it with the data of a single blood donor to ensure that only the morphological features driven by the agonists contributed to the development of the iPAC (Figure 1C; Transparent Methods). In comparison, we measured the platelet samples that were prepared under the same procedure using a conventional flow cytometer (Cytomics FC500, Beckman Coulter). As shown in Figure 2, the flow cytometer was not capable of differentiating them as indicated by their significant overlap (Table S1; Transparent Methods).

**Figure 1.**
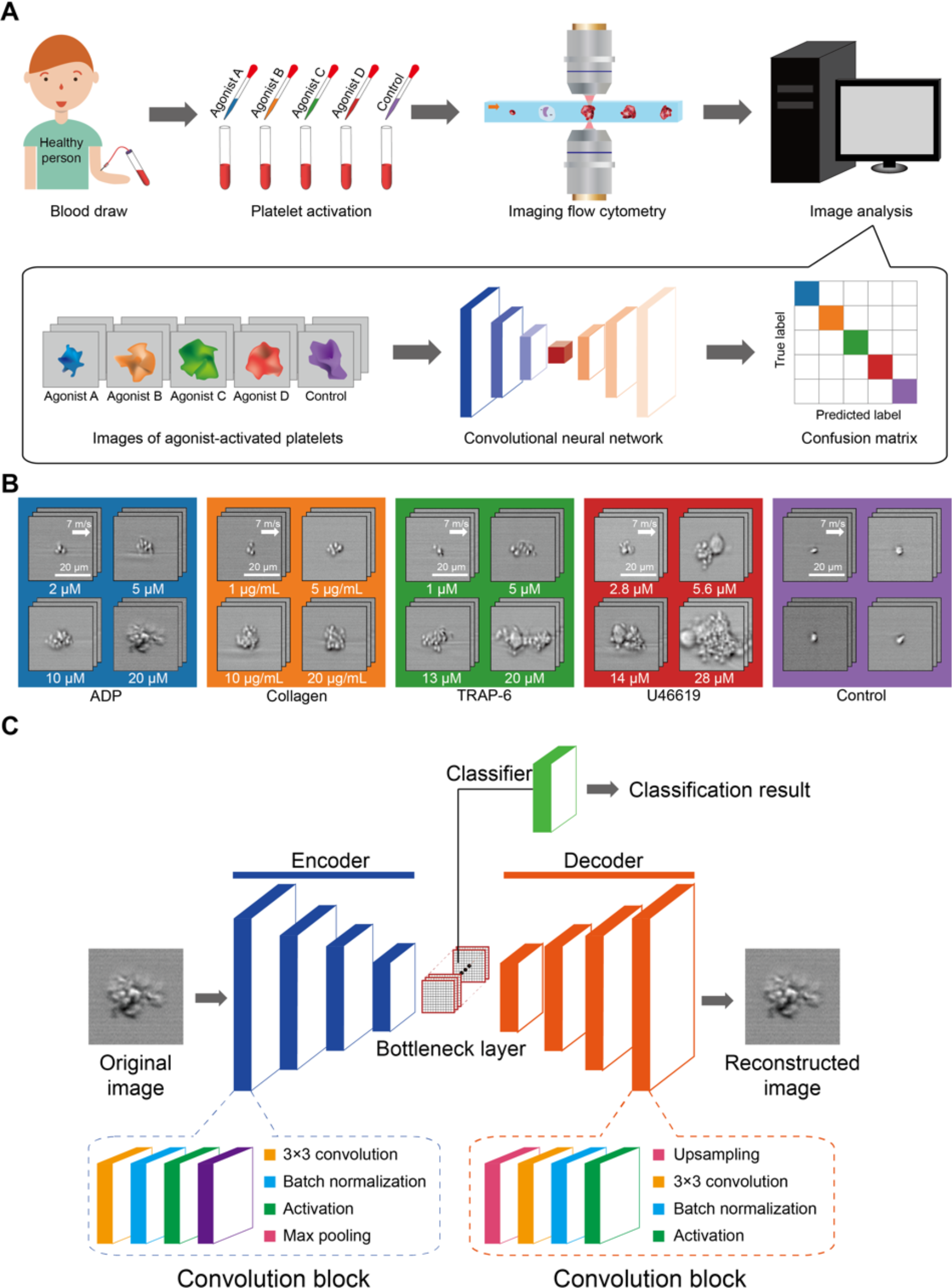
Development of the iPAC. (A) Procedure for developing the iPAC. (B) Images of the agonist-activated platelet aggregates and single platelets (negative control). (C) Structure of the CNN with an encoder-decoder architecture used for the development of the iPAC.

**Figure 2.**
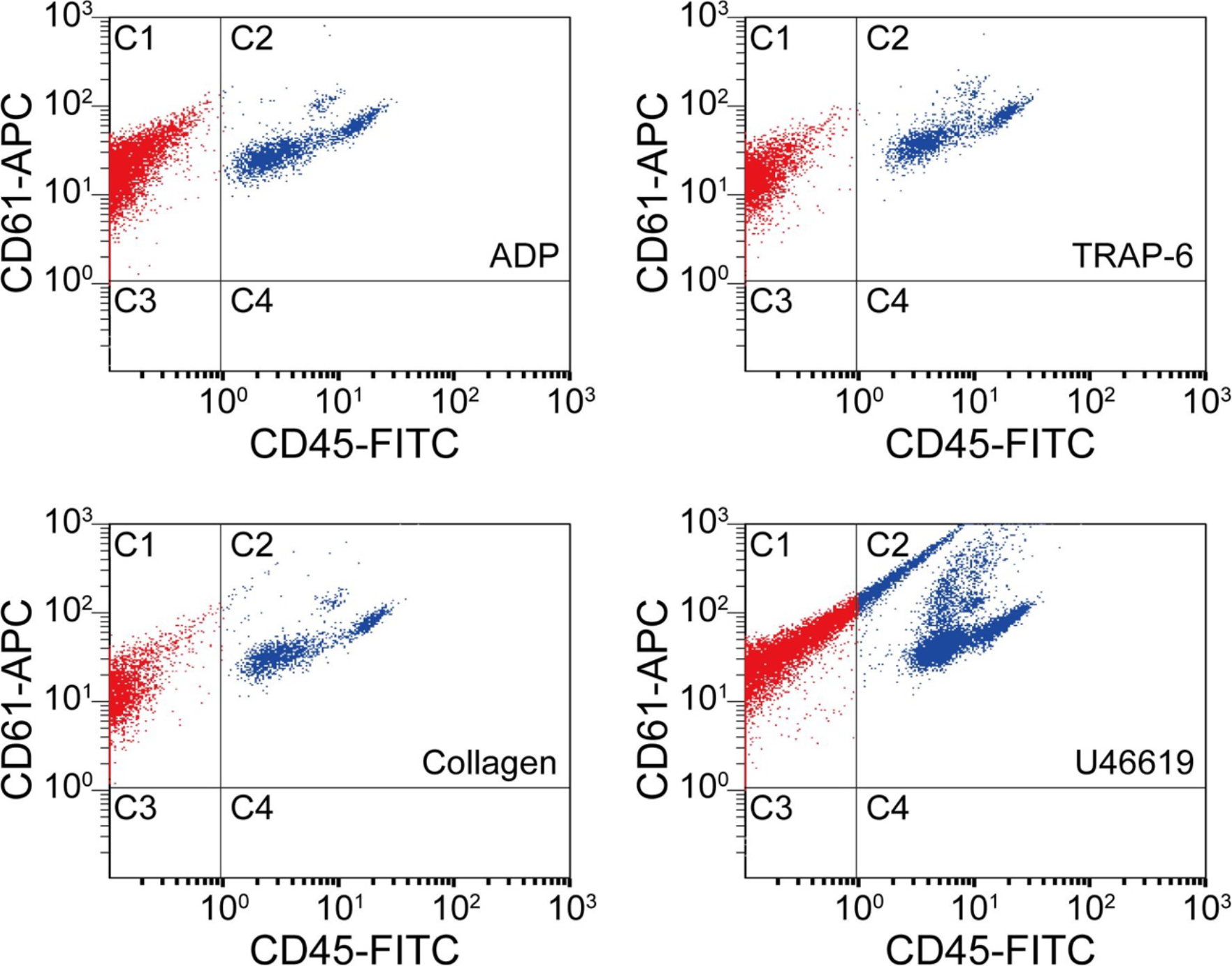
Scatter plots of agonist-activated platelets analyzed by a conventional flow cytometer. C1: single platelets and platelet-platelet aggregates. C2: leucocytes and platelet-leucocyte aggregates.

### Demonstration of the iPAC

The iPAC is manifested as a confusion matrix with each row representing the examples in a predicted class and each column representing the examples in an actual or true class. As shown in Figure 3A, most of the images were classified into the correct groups in the diagonal line of the confusion matrix. Large separations between the different platelet sample portions in Figure 3B that visualizes the bottleneck layer in the CNN indicate the first CNN model’s ability to discriminate various types of agonist-activated platelet aggregates and negative control. The negative control shows the highest classification accuracy, indicating that large morphological changes were made to the activated platelets. The U46619-treated blood sample portion shows the second highest classification accuracy of all the blood sample portions, indicating that the morphological changes caused by the agonist are very different from those caused by the other agonists. Many platelet-leukocyte aggregates were identified in the U46619-treated sample portion, but few in the other blood sample portions (Figure 2). This may be because U46619 acted as a thromboxane A2 (TXA_2_) receptor agonist, which activated endothelial TXA_2_ receptors, promoting the expression of adhesion molecules, and thus favored adhesion and infiltration of leukocytes (Michelson, 2012; George, 2000). The low classification accuracy values of the ADP-, collagen-, and TRAP-6-treated blood sample portions are presumably due to the fact that these agonists partially share similar mechanisms in forming platelets aggregates (Michelson, 2012; George, 2000; Michelson, 2003; Harrison, 2005; Li et al., 2000). For example, since platelets also release ADP themselves during activation (Michelson, 2012; George, 2000; Michelson, 2003; Harrison, 2005), platelet aggregates produced by other agonists may also share similar morphological features as ADP-activated platelet aggregates. To demonstrate the reproducibility of the iPAC, we tested it with an independent dataset (a total of 25,000 images of all event types), which was performed under the same conditions as shown in Figure 1A. The contribution values over all the agonists are in good agreement with the values in the diagonal elements of the confusion matrix (Figure 3C), which validates the reliability of the iPAC.

**Figure 3.**
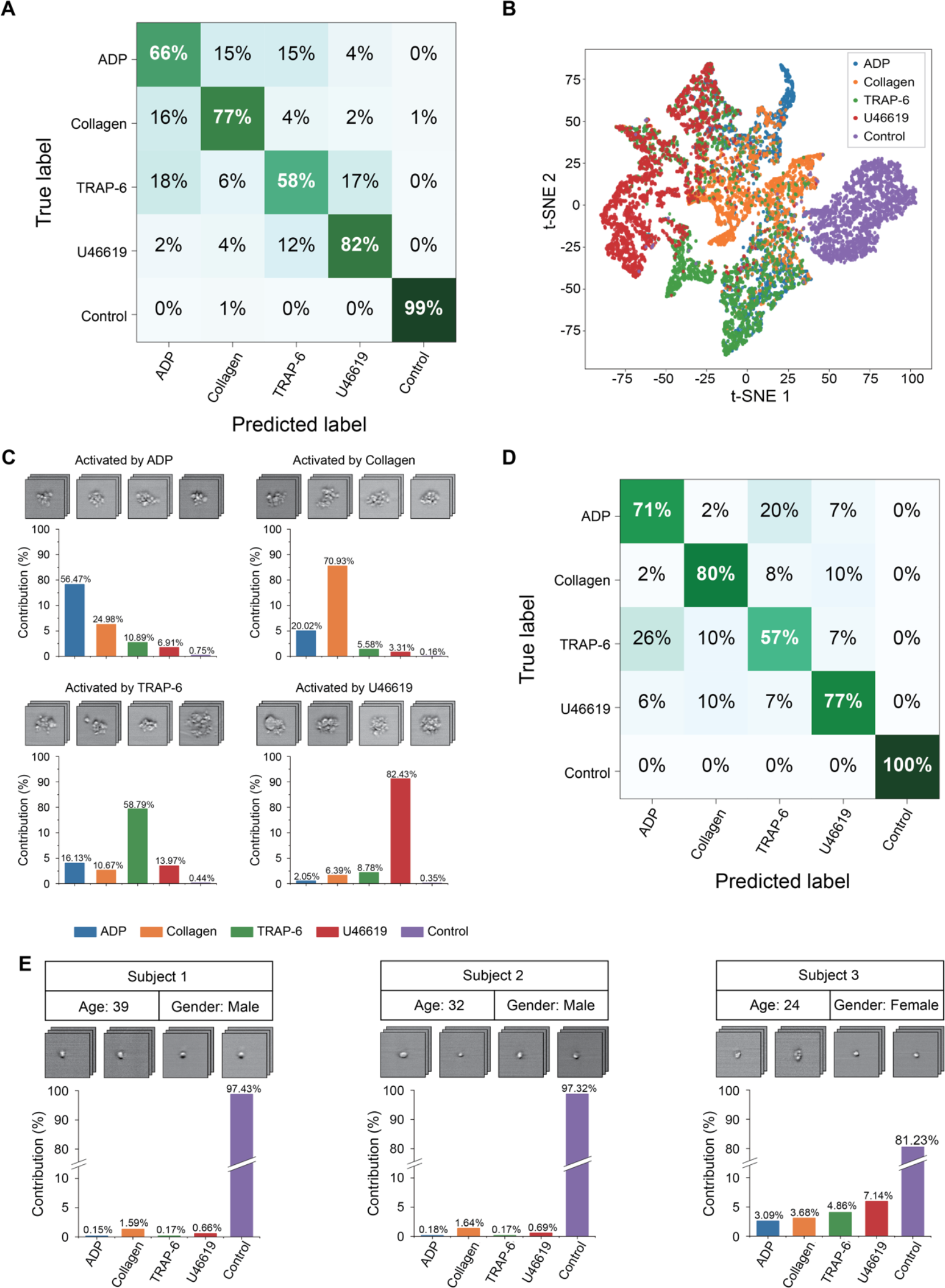
Demonstration of the iPAC. (A) Confusion matrix as a manifestation of the iPAC. (B) t-SNE plot of the agonist-activated platelet aggregates and single platelets (negative control). (C) Validation of the reproducibility of the iPAC. (D) Confusion matrix of the CNN model trained with the images of platelet aggregates activated by different concentrations of agonists. (E) iPAC-based diagnosis of platelets from three healthy human subjects.

The iPAC’s ability to classify platelet aggregates by agonist type in a concentration-independent manner is indicated by the confusion matrix shown in Figure 3D with an average diagonal element value of 77%. The results also reveal the existence of the unique morphological features related to each agonist type, which is promising for potential application to diagnosis of thrombotic disorders by tracing back to the leading factors of platelet aggregation. In addition, from a viewpoint of potential clinical applications, while the conventional assays can only evaluate platelet aggregability qualitatively, the iPAC can quantify it with the resolving power to identify the contribution of each agonist type to it. However, it can be recognized from the image library (Figure 1B) that U46619-activated platelet aggregates have relatively larger size than those in the other sample portions, which may be captured as a type of morphological features by the CNN, leading to the high classification accuracy of the U46619-activated samples.

To demonstrate the diagnostic utility of the iPAC, we applied it to blood samples of three healthy human subjects to predict the contribution of each agonist type to platelet aggregates (if any) in the samples (Figure 3E). The blood samples were prepared by following the same procedure as shown in Figure 1A except for the step of adding agonists (with 5,000 images of events in each blood sample). Only 2% of the total population of platelets in the first two blood samples were identified as aggregates, whereas the third blood sample was found to contain a relatively large concentration of platelet aggregates mainly activated by TRAP-6 and U46619. The iPAC’s diagnostic ability to obtain this type of information is an effective tool for studying and elucidating the mechanism of platelet aggregation and holds promise for clinical diagnostics, pharmacometrics, and therapeutics, although the iPAC needs more training with a wide spectrum of diseases and medical conditions for the purpose. For example, the iPAC may provide an important clue to the choice of drugs (e.g., aspirin or thienopyridines) for antiplatelet therapy (Mauri et al., 2014; Roe et al., 2012), the gold standard of the treatment and prevention of atherothrombosis (e.g., myocardial infarction, stroke), in that aspirin inhibits the formation of TXA_2_ whose stable analogue is U46619 while thienopyridines exert an antiplatelet effect by blocking the ADP receptor P2Y_12_. Furthermore, the iPAC may be able to identify TRAP-6-activated platelet aggregates in the bloodstream of patients with deep vein thrombosis (since TRAP-6 interacts with the receptor of thrombin) and suggest that they come from the venous side.

## DISCUSSION

The information about the driving factors behind the formation of platelet aggregates is expected to lead to a better understanding of the underlying mechanisms of platelet aggregation and, thereby, open a window on an entirely new class of clinical diagnostics and therapeutics. For example, antiplatelet therapy is the gold standard of the treatment and prevention of atherothrombosis (e.g., myocardial infarction and stroke) for which aspirin and thienopyridines (e.g., prasugrel and clopidogrel) are primarily used as antiplatelet drugs worldwide (Mauri et al., 2014; Roe et al., 2012). Aspirin inhibits the formation of TXA_2_ whose stable analogue is U46619, whereas thienopyridines exert an antiplatelet effect by blocking the ADP receptor P2Y_12_. Accordingly, the ability to identify the type of platelet aggregates in the blood stream may provide an important clue to the choice of a drug for antiplatelet therapy. Furthermore, deep vein thrombosis (DVT) is a blood clot that normally occurs in a deep vein where coagulation activation plays an important role. Since TRAP-6 interacts with the receptor of thrombin (i.e., the product of the coagulation cascade), the ability to identify TRAP-6-activated platelet aggregates in the blood stream may suggest that aggregates come from the venous side. Therefore, the iPAC may pave the way for introducing a novel laboratory testing technique for the management of pathological thrombosis such as atherothrombosis and DVT although further basic and clinical studies are needed.

The relation between platelet activation signaling pathways and the formation of platelet aggregates has been extensively studied (Li et al., 2010; Michelson, 2012; Brass, 2013). It is known that agonists activate platelets in a selective manner via specific receptors, which is followed by a variety of downstream signaling events (Li et al., 2010). For example, collagen interacts with the immune-like receptor glycoprotein VI, which signals through an immunoreceptor tyrosine-based activation motif and activates the tyrosine phosphorylation pathway (Michelson, 2012; Li et al., 2010) In contrast, soluble agonists such as TRAP-6, U46619, and ADP interact with G protein-coupled receptors (Michelson, 2012; Brass, 2003). Furthermore, each soluble agonist couples with a specific type of G protein, which leads to different aggregation mechanisms (Rivera et al., 2009) and thus suggests different underlying mechanisms for expressing different morphological features on platelet aggregates. It is challenging, but is expected to be intriguing to study and elucidate the mechanisms for a further understanding of the biology of platelets.

## TRANSPARENT METHODS

- Detailed methods are provided in the online version of this paper and include the following:
- KEY RESOURCES TABLE
- CONTACT FOR REAGENT AND RESOURCE SHARING
  ○ Blood samples for detection of platelet aggregates
- EXPERIMENTAL MODEL AND SUBJECT DETAILS
- METHOD DETAILS
  ○ Microfluidic chip fabrication
  ○ Optofluidic time-stretch microscopy
  ○ Evaluation of agonist-activated platelets by conventional flow cytometry
- QUANTIFICATION AND STATISTICAL ANALYSIS
  ○ Convolutional neural network
- DATA AND SOFTWARE SHARING

## ACKNOWLEDGEMENTS

This work was supported by the ImPACT Program, JSPS C2C Program, White Rock Foundation, Nakatani Foundation, and University of Tokyo’s Center for Nano Lithography.

## AUTHOR CONTRIBUTIONS

C.L., Y.Y., and K.G. conceived the original idea. Y.Z. and A.Y. performed the optical and biological experiments. Y.Z. and C.J.H. analyzed the results. H.K., Y.W., S.Y., C.W.S., and Y.Y. provided assistance to the experiments and manuscript writing. Y.Z., C.L., and K.G. wrote the manuscript. C.L. supervised the experiments and data analysis. K.G. supervised the team. All authors participated in revising the manuscript.

## DECLRARATION OF INTERESTS

K.G. is a shareholder of two cell analysis startups.

