## Supplementary information for "Classification of platelet aggregates by agonist type"

#### Supplemental Information

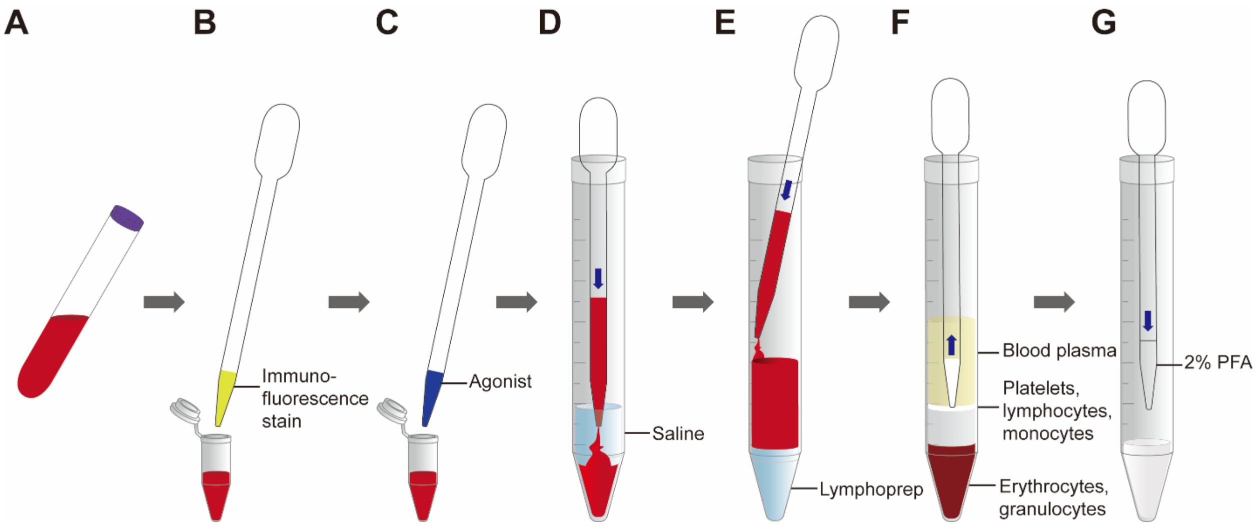


**Figure S1. Detailed Procedure for Sample Preparation, Related to Figure 1**

(A) Blood is drawn into a blood collection tube containing citric acid.

(B) Platelets are labeled by immunofluorescence.

(C) An agonist is added to the tube.

(D) The agonist-activated blood sample is diluted with saline.

(E) The sample is loaded on top of the Lymphoprep followed by centrifugation.

(F) Enriched platelet aggregates are collected from the mononuclear layer.

(G) Enriched platelet aggregates are fixed by 2% paraformaldehyde.

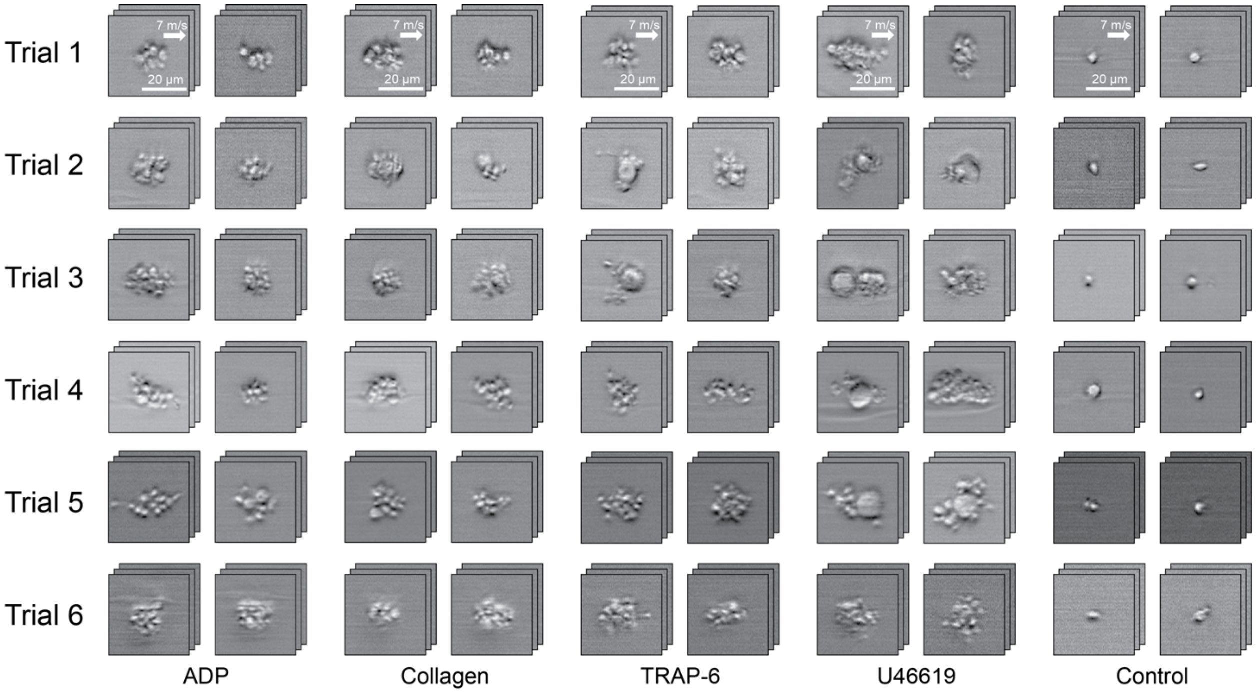


**Figure S2. Extend Image Library from Six Experimental Trials, Related to Figure 1**

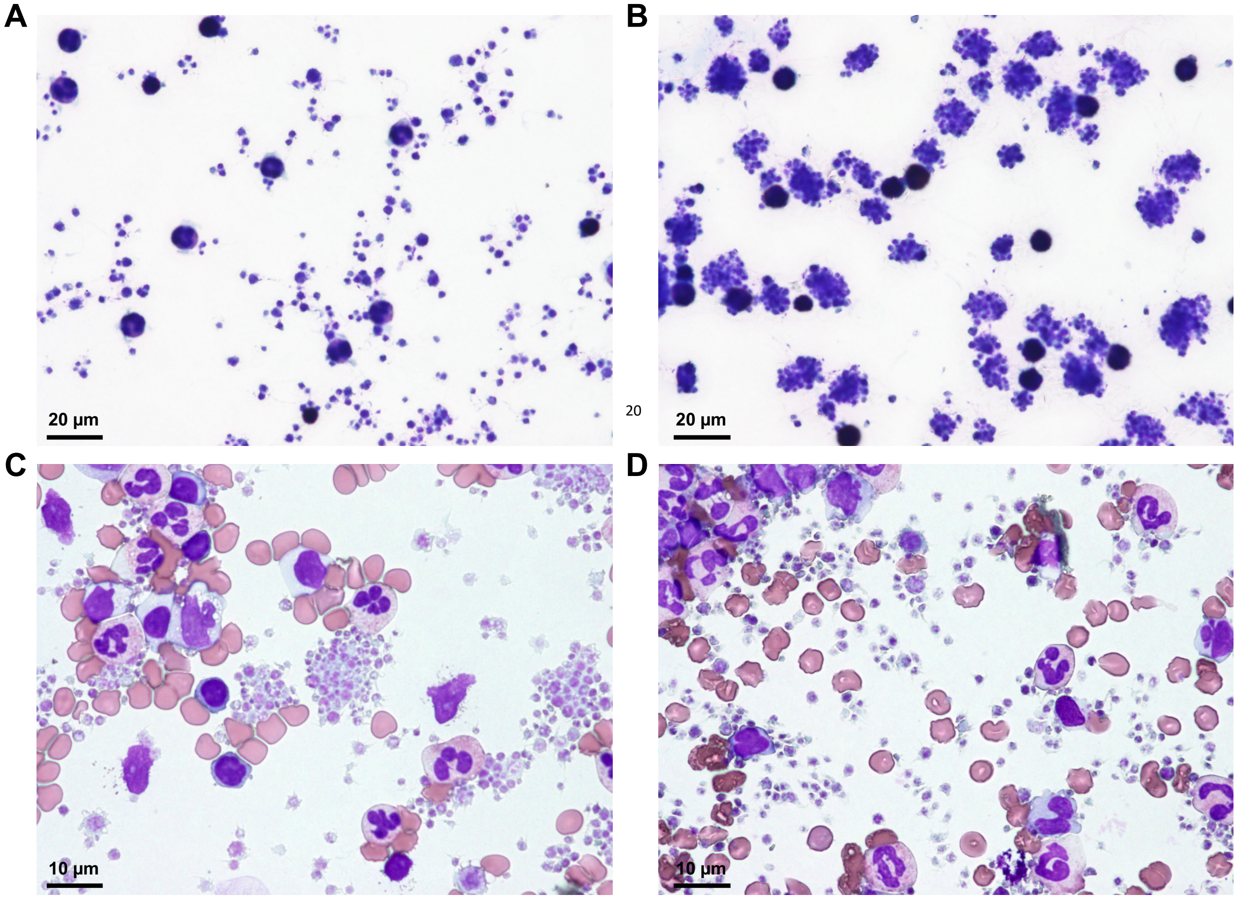


**Figure S3. Comparing Different Procedures for Sample Preparation, Related to Figure 1**

(A) Microscopic image of a platelet sample prepared by pipetting.

(B) Microscopic image of a platelet sample prepared by vortexing.

(C) Microscopic image of a platelet sample without fixation at 0 hr.

(D) Microscopic image of a platelet sample without fixation at 3 hr.

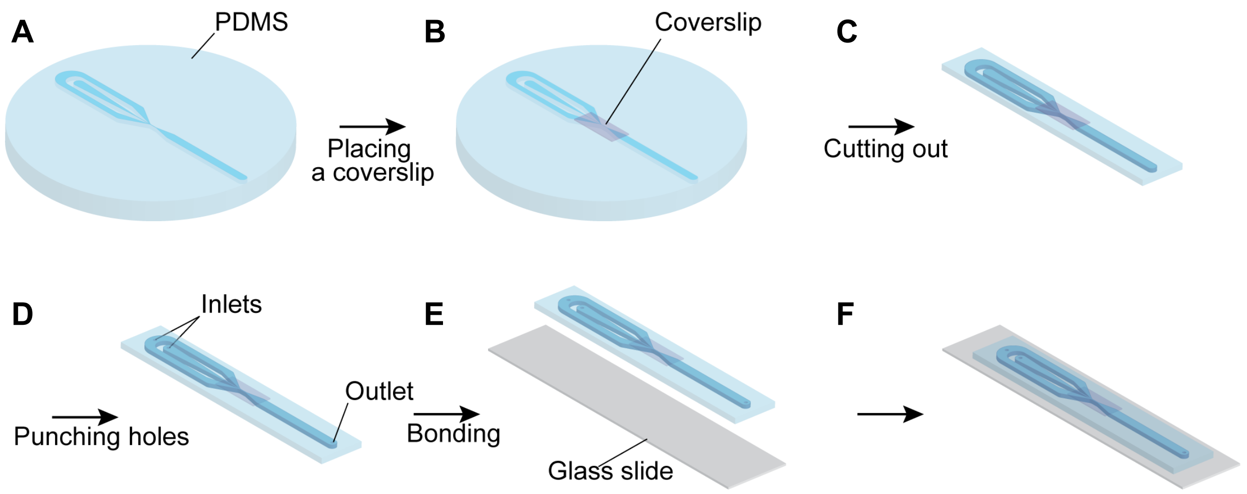


**Figure S4. Fabrication of the Microfluidic Chip, Related to Figure 1**

(A) PDMS is filled on the mold of the microfluidic channel.

(B) The small piece of the coverslip is placed above the observation area on the PDMS.

(C) The PDMS is cut out into a proper size.

(D) The inlets and outlet holes are opened.

(E) The PDMS and glass slide are bonded using a plasma cleaner.

(F) The microfluidic chip is completed and ready for use.

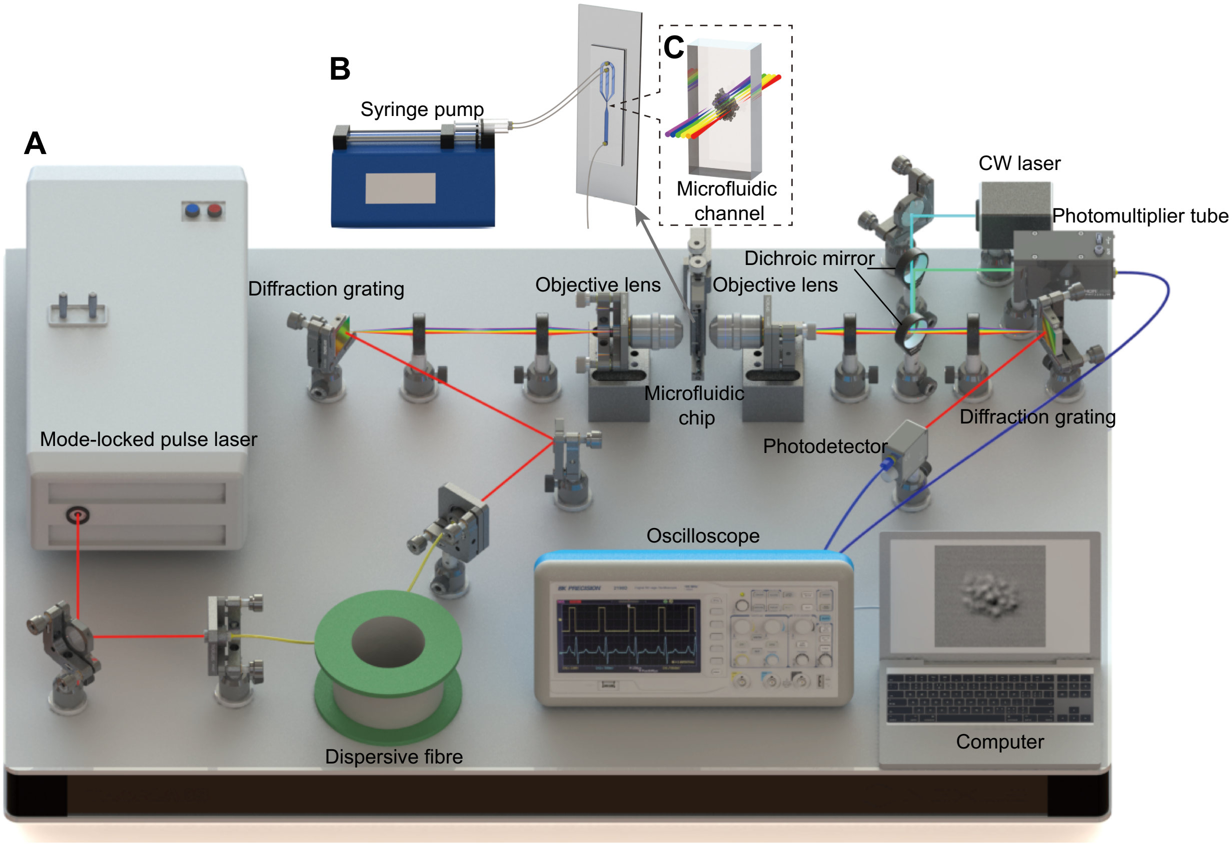


**Figure S5. Schematic of the Optofluidic Time-stretch Microscope, Related to Figure 1**

(A) Overall schematic.

(B) Enlarged schematic of the microfluidic chip.

(C) Further enlarged schematic of the microfluidic channel.

**Table S1. Statistical analysis of agonist-activated platelets by conventional flow cytometry, Related to Figure 2**

The population and ratio of dots in each area are shown in the table for all the four types of agonists. C1 areas indicate single platelets and platelet-platelet aggregates. C2 areas indicate platelet-leucocyte aggregates. C3 areas indicate blood cells other than platelets and leucocytes. C4 areas indicate leucocytes.

| Agonist | Area | Population | Ratio |
|---|---|---|---|
| ADP | C1 | 45260 | 90.52% |
| | C2 | 4708 | 9.42% |
| | C3 | 32 | 0.06% |
| | C4 | 0 | 0.00% |
| Collagen | C1 | 46672 | 93.34% |
| | C2 | 3303 | 6.61% |
| | C3 | 25 | 0.05% |
| | C4 | 0 | 0.00% |
| TRAP-6 | C1 | 46297 | 92.59% |
| | C2 | 3689 | 7.38% |
| | C3 | 14 | 0.03% |
| | C4 | 0 | 0.00% |
| U46619 | Ç1 | 34030 | 68.06% |
| | C2 | 15924 | 31.85% |
| | C3 | 46 | 0.09% |
| | C4 | 0 | 0.00% |

### Transparent Methods

#### KEY RESOURCES TABLE

#### CONTACT FOR REAGENT AND RESOURCE SHARING

#### EXPERIMENTAL MODEL AND SUBJECT DETAILS

##### Blood samples for detection of platelet aggregates

The detailed procedure of the sample preparation is shown in Figure S1, where platelets and platelet aggregates were enriched from whole blood by the density-gradient centrifugation to maximize the detection efficiency (Beakke, 1951). Specifically, blood samples were obtained from a healthy person with citric acid as the anticoagulant, which did not dissociate platelet aggregates (Figure S1A). Platelets were immunofluorescently labeled by adding 20-μL PE anti-human CD61 (BioLegend, 336405) to the blood samples to ensure that platelets would be detected in all images (Figure S1B). For each agonist type, 500-μL blood was incubated with 50-μL agonist solution, which contained 20-μM ADP (BioMed, AP-200-422), 10-μg/mL Collagen (BioMed, AG005K-CS), 13-μM TRAP-6 (H2936.0005, BACHEM), or 14-μM U46619 (Cayman Chemical, 16450), for 10 min (Figure S1C). The labeled, activated platelets were then diluted using 5-mL saline (Figure S1D). Next, the platelets were isolated by using Lymphoprep (STEMCELLS, ST07851), a density-gradient medium, using the protocol provided by the vendor. Specifically, the diluted blood was added on top of the Lymphoprep and then centrifuged at 800 g for 20 min (Figure S1E). After the centrifugation, 1 mL of the sample was taken from the mononuclear layer, to which 1 mL of 2% paraformaldehyde (Wako, 163-20145) was added for fixation (Lanier and Warner, 1981) (Figures S1F and S1G). The operation of the fixation was performed at 4°C for 30 min while other operations were performed at 25°C room temperature. As shown in Figure S2, we first compared several procedures of preparing blood samples, but most of the procedures either left a large amount of non-target blood cells in the sample, thus decreasing the iPAC's detection efficiency, or dismantled the agonist-activated platelet aggregates. The current procedure is advantageous over the procedures in preserving the morphology of platelet aggregates while eliminating non-target blood cells. This study was approved by the Institutional Ethics Committee in the School of Medicine at the University of Tokyo [no. 11049-(6)]. Written informed consents were obtained from the blood donors.

#### METHOD DETAILS

**Microfluidic chip fabrication**

The microfluidic chip was fabricated using standard photolithographic methods (Whitesides et al., 2001). A designed pattern of the microfluidic channel was drawn using AutoCAD (Autodesk) and printed on a film mask (UnnoGiken). Negative photoresist (KMPR 1035, MicroChem) was spin-coated on a silicon wafer and heated at 100°C for 10 min. Then, the silicon wafer, covered with the film mask, was exposed to ultraviolet (UV) light followed by hard baking at 100°C for 5 min and developed using SU-8 developer (MicroChem). After washing with isopropyl alcohol and water, the silicon wafer was heated at 150°C for 15 min. The negative photoresist mold on the silicon wafer was fixed in a Petri dish and then filled with polydimethylsiloxane (PDMS, Dow Corning) in which PDMS base and curing reagent were mixed at a ratio of 10:1 (Figure S4A). PDMS was heated at 80°C for 15 min, and then a small piece of coverslip was placed on PDMS right above the observation area of the microfluidic channel. This step improved the mechanical strength of PDMS so that the channel (Figure S4B) was able to resist the pressure inside the channel without deformation. After another heating for more than 1 h, the PDMS layer was cut into a small piece so that it could fit in the size of a glass slide (Figure S4C). The inlets and outlet were punched by a 25G needle (Figure S4D). To form permanent bonding between the PDMS channel and the glass slide, both the PDMS device and the glass slide were treated with a plasma cleaner (Harrick Plasma) (Figure S4E). The dimensions of the microchannel in the observation area are about 80 μm in width and 40 μm in height (Figure S4F).

**Optofluidic time-stretch microscopy**

The optofluidic time-stretch microscope (Lei et al., 2016) is schematically shown in Figure S5. A Ti:Sapphire mode-locked femtosecond pulse laser with a center wavelength, bandwidth, and pulse repetition rate of 780 nm, 40 nm, and 75 MHz, respectively, was used as an optical source. Each laser pulse was first stretched temporally by a single-mode dispersive fiber with a group-velocity dispersion of -240 ps/nm (Nufern 630-HP) and then dispersed spatially by the first diffraction grating with a groove density of 1,200 lines/mm. The stretched laser pulse was focused by the first objective lens (Olympus, 40×, NA 0.6) onto a flowing cell in the microfluidic channel. The pulse that contained the spatial profile of the cell on its spectrum was collected by the second objective lens and spatially recombined by the second diffraction grating, followed by photodetection with a high-speed photodetector (New Focus 1580-B) with a detection bandwidth of 12 GHz. To ensure imaging of platelet-related events (i.e., single platelets, platelet-platelet aggregates, platelet-leukocyte aggregates), fluorescence detection was used in conjunction with the optofluidic time-stretch microscope. A 488-nm continuous-wave laser was used to detect CD61 fluorescence signals with a photomultiplier tube (Hamamatsu H10723-01MOD). Only the image signals associated with CD61 fluorescence signals were collected. The image-encoded pulse and fluorescence signal were digitized using a high-speed oscilloscope (Tektronix DPO 71604B) with a detection bandwidth of 16 GHz and a sampling rate of 50 GS/s. Pulses were repeated by the mode-locked pulse laser at 75 MHz so that image-encoded pulses detected by the photodetector were digitally stacked to form 2D images using MATLAB R2018b (MathWorks). The pulse intensity profile (usually Gaussian-shaped with ripples) was normalized to obtain a flat background. Also, the images were cropped into 160×160 pixels to obtain only the cell-contained parts for further analysis.

**Evaluation of agonist-activated platelets by conventional flow cytometry**

We analyzed agonist-activated platelets with a conventional flow cytometer (Cytomics FC500, Beckman Coulter) that can count and analyze large cell populations via scattering and fluorescence measurements with high throughput. Blood samples were processed using the same procedure as for optofluidic time-stretch microscopy, but labeled with anti-CD61-APC and anti-CD45-FITC antibodies (Beckman Coulter) for detecting white blood cells and platelets, respectively. To only detect single platelets and platelets aggregates, gating of cellular size and granularity was applied to the light scatter plots. As shown in Figure 2, the C1 areas, which correspond to CD61-APC positive and CD45-FITC negative, show events associated with single platelets and platelet-platelet aggregates. The C2 areas, which correspond to CD61-APC/CD45-FITC double positive, show events associated with platelet-leukocyte aggregates. The C3 areas (CD61-APC/CD45-FITC double negative) and C4 areas (CD61-APC negative and CD45-FITC positive) correspond to events which did not contain any platelets.

#### QUANTIFICATION AND STATISTICAL ANALYSIS

**Convolutional neural network**

The details of the CNN with the encoder-decoder architecture are as follows. The encoder was used to extract morphological features of platelet aggregates, while the decoder was used to recover the platelet aggregate images from the morphological features. This two-stage structure forced the encoder to extract features from the cells instead of the background or noise, which helped enhance the reliability and accuracy of classification. The images were normalized to 0-mean and divided into training, validation, and test sets at a ratio of 3:1:1. The CNN classifier was trained on the training set. The validation loss was calculated with the validation dataset at each epoch to monitor the learning process. The learning rate was reduced when the validation loss stopped descending for more than 3 epochs until it reached $1\times10^{-8}$. The training was ceased when there was no more decrease in the validation loss for more than 6 epochs. After the training ended, the test set was processed to calculate the final classification accuracy for each agonist type. The CNN classifier was implemented on Keras (Chollet, 2015) with the Tensorflow (Abadi et al., 2016) backbone. The training of the CNN classifier was optimized by Adam with an initial learning rate of 0.001.

#### DATA AND SOFTWARE AVAILABILITY

The image data obtained by the optofluidic time-stretch microscope and the code used to analyze the images and develop the confusion matrix are available upon request.
